# Enriched environment requires remodeling of hippocampal perineuronal nets to trigger memory improvement in Alzheimer’s mouse model

**DOI:** 10.1101/2024.10.24.619979

**Authors:** Guillaume Bouisset, Fanny J. Tixier, Tatiana Dupak, Camille Lejards, Laure Verret

**Author notes:** Instituto de Neurociencias CSIC-UMH, San Juan de Alicante, Spain.

## Abstract

Alzheimer’s disease (AD) is a major neurodegenerative disorder influenced by both genetic and environmental factors. Engaging in mentally stimulating activities is believed to reduce cognitive decline by establishing a cognitive reserve, though the underlying neurobiological mechanisms remain elusive. In this study, we explore the role of parvalbumin-expressing inhibitory neurons (PV) and their associated perineuronal nets (PNN) in cognitive deficits observed in AD. Using the Tg2576 mouse model, we demonstrate that 10 days of exposure to an enriched environment (EE) significantly restores spatial and social memory, accompanied by an increase in PV and PV/PNN cell populations in the hippocampus. Notably, preventing PV/PNN remodeling in the CA1 region during EE abolishes the spatial memory improvements, whereas localized neuregulin-1 (NRG1) injections induce PV/PNN remodeling and restore memory function. These findings suggest that hippocampal PV/PNN remodeling is essential for the cognitive benefits of EE in AD, highlighting this neuronal population as a critical substrate for cognitive reserve. This study provides new insights into the mechanisms by which environmental factors may mitigate cognitive decline in AD, offering potential avenues for therapeutic interventions.

**Highlights:** 10-day environmental enrichment increases PV+ and PV+/PNN+ in the hippocampus of Tg2576 mouse model of AD

Blocking PV/PNN remodeling during EE prevents memory recovery in Tg2576 mice

Enhancing PV/PNN remodeling with NRG1 leads to restored memory in Tg2576 mice

PV/PNN remodeling in area CA1 affects spatial memory, while in CA2, it impacts social memory

## Introduction

Alzheimer’s disease (AD) is currently the most prevalent neurodegenerative disease among the aging population worldwide. This progressive brain condition is associated with inevitable impairment of cognitive functions, including memory. Despite the myriads of research efforts, including preclinical studies and cIinical trials aimed at designing disease-modifying and/or cognitive enhancing drugs, no effective treatment is presently accessible for patients^1^.

Additionally, it is well established that the susceptibility to developing dementia in AD is influenced by environmental factors and level of education ^2,3^, and that certain lifestyle changes may potentially reduce the risk of cognitive decline and dementia ^4^. Indeed, individuals leading a mentally stimulating lifestyle could harness their passive, pre-existing cognitive processes to mitigate the adverse impact of the pathology ^5,6^. Among the contributors to this cognitive reserve, a higher level of education ^7^ or occupational complexity ^8^ could potentially enhance cognitive status and reduce the likelihood of progressing to dementia ^9^. Comprehending how these factors modify the brain to provide better coping mechanisms against pathological insults and to delay the clinical expression of the disease could be pivotal in the development of effective non-pharmacological interventions, as well as novel drug-design strategies.

Learning and memory retrieval necessitate a high level of information processing, that is supported by coordinated activity of neuronal assemblies through balanced excitatory and inhibitory transmission ^10^. Over the last two decades, mounting evidence has demonstrated that both AD patients and mouse models exhibit an imbalance between excitation and inhibition (E/I), leading to spontaneous epileptic-like activities and altered brain oscillations. These aberrant brain activities substantially contribute to, if not directly underlie, the observed cognitive deficits ^11–13^. In line with their crucial role in maintaining the E/I balance in the brain ^14^, we and others have established a causal link between the impaired function of parvalbumin-positive (PV) GABAergic neurons and both abnormal brain network activity and cognitive impairments in AD mouse models and patients ^11,15,16^. In the adult–hence mature–brain, PV cells are often associated with the perineuronal net (PNN), a dense extracellular matrix organized into a lattice-like structure ^17^. PNNs develop around PV cells in an experience-dependent manner during the post-natal maturation of the cortex and the hippocampus ^18^. It has been proposed that their presence is crucial for maintaining existing synapses, as well as preventing the formation of new ones, on mature PV cells. Therefore, PNNs stabilize excitatory inputs onto inhibitory PV neurons; their presence is associated with an increased firing rate ^19,20^ and PV protein level ^21^. As they form concomitantly with the closure of critical periods of plasticity, PNN presence is often considered to reduced plasticity ^18,22^. However, PNN formation can occur throughout life, allowing consolidation of synaptic inputs onto PV cells, ensuring maintenance of neural networks ^19^, and protecting memories from erasure ^23^. Furthermore, the post-natal maturation of PNN around PV cells in hippocampal CA1 and CA2 areas is a prerequisite for the emergence of the ability to form episodic-like (contextual and social) memory in adult mice ^24,25^.

In agreement with some observations in the human brain ^26^, we recently showed that mice of the Tg2576 mouse model of AD have decreased PV expression as young as 3 months, *i.e.*, before amyloid-beta peptides (Aβ) overproduction ^27^, which is comparable to what has been observed in a previous report ^28^. As PNN presence appears to be necessary for memory formation and recall ^24,25^, as well as network stabilization, its absence in the hippocampus of the Tg2576 mouse model of AD even during the presymptomatic stage ^27,29^, may contribute to memory dysfunction. In line with this, strategies aiming at reforming PNN around PV cells of Tg2576 mice result in improvement of their memory performance. Specifically, a 10-week stay in an enriched environment (EE) during early adulthood restores PV expression and PNNs in the hippocampus of Tg2576 mice ^27^, and has long-lasting (at least 8 months) beneficial effects on their cognitive performance ^30^. Interestingly, in rodent models, transient stay in an EE has been widely used to simulate a complex social environment, as well as sensory and cognitive stimulations, known to provide cognitive reserve in humans ^31,32^. Taking into account these observations, we hypothesize that the cognitive benefits induced by EE in AD individuals are sustained by the remodeling of PV interneurons and their PNN. We thus prevented the remodeling of PV cells in a specific hippocampal area (CA1) throughout the period of enriched housing in Tg2576 mice, with the bacterial enzyme Chondroitinase-ABC (ChABC). We consequently observed no improvement of spatial memory, linked to CA1 function, specifically in the Tg2576 mice where PV/PNN remodeling was prevented during EE, whereas their social memory performance was restored. Therefore, we demonstrate that the change in PV cell network could sustain the long-lasting beneficial impact of EE in AD mice.

## Results

### ChABC injection in CA1 prevents local PV cell remodeling during EE

We used Chondroitinase-ABC (ChABC) to digest PNNs and thereby prevent the remodeling of PV-positive (PV+) neurons during a 10-day period. ChABC is a bacterial enzyme known to digest PNNs around PV+ cells, and previous studies have shown that PNNs remain absent for up to 2 weeks following a single injection ^33,34^. Consistent with this, we observed that ChABC injection into the CA1 area of non-transgenic (NTg) mice housed in standard conditions (SH) led to a decrease in the number of detectable PV+/PNN+ (373.30 [0.74 to 745.90]) and PV+ cells (536.3 [52.76 to 1021]) 10 days after injection (**Fig.1A-C,** two-way ANOVA, PV+/PNN+, interaction: F(1,23) = 4.328; p = 0.0488, Tukey’s post-hoc test: NTg-PBS-SH *vs* NTg-ChABC-SH, p = 0.0494; two-way ANOVA, PV+, interaction: F(1,23) = 5.651; p = 0.0261,Tukey’s post-hoc test: NTg-PBS-SH *vs* NTg-ChABC-SH, p = 0.0260). Interestingly, this led to a deficit in PV and PNN cell numbers comparable to that observed in CA1 area of Tg2576 mice without ChABC (**Fig.1B**). As a control for the accuracy of injection in CA1, the numbers of PV+/PNN+ and PV+ cells in the CA2 area of NTg mice were not affected by the ChABC injection in CA1 (**Fig.S1A-B**).

**Figure 1:**
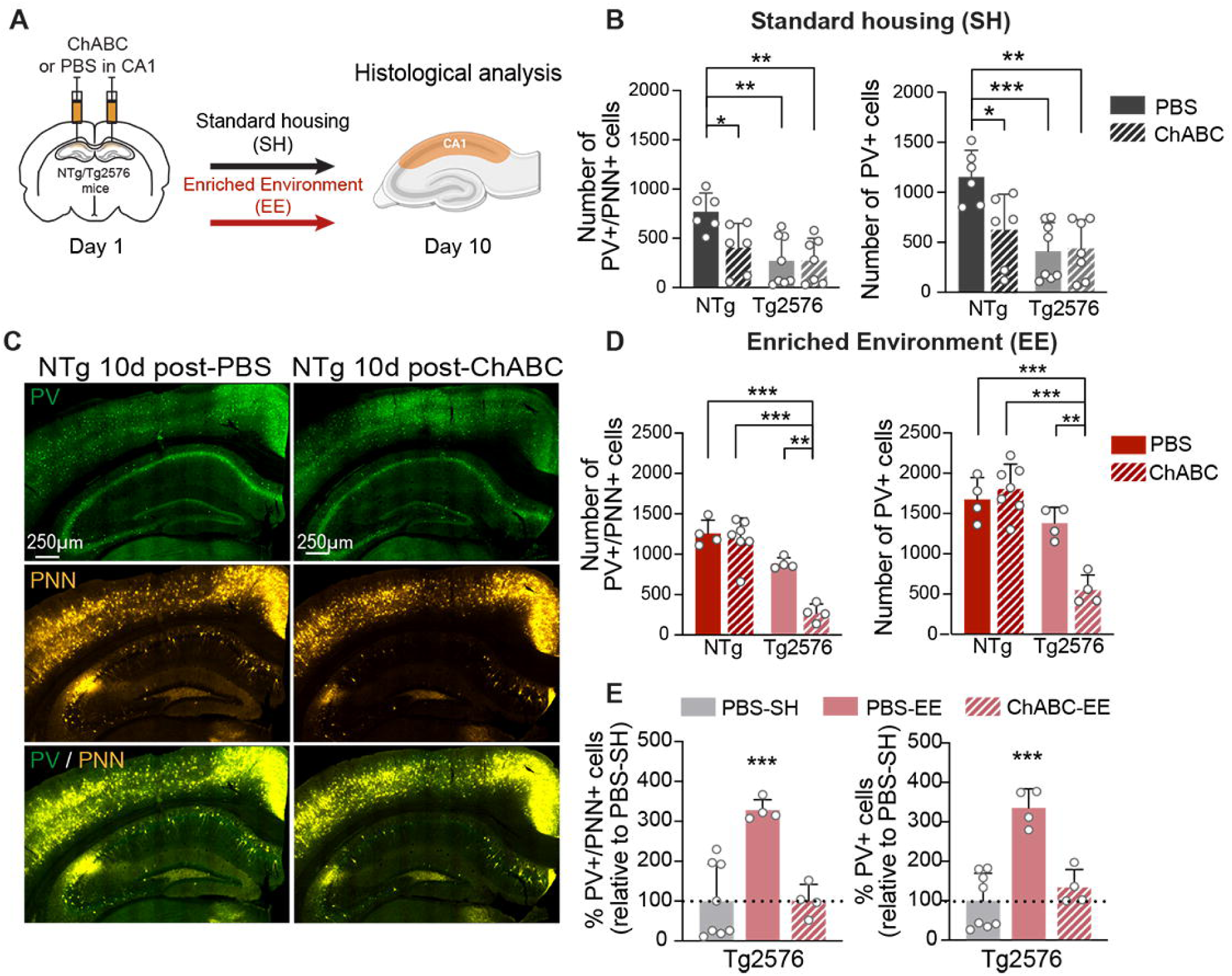
ChABC Injection in CA1 Prevents Local PV Cell Remodeling During ***EE.* (A)** Experimental timeline: ChABC or PBS was injected into the CA1 area of NTg and Tg2576 mice, which were then housed in standard (SH) or enriched (EE) conditions. Mice were sacrificed for PV and PNN staining 10 days post-injection. **(B)** In standard housing conditions (SH), ChABC injection into the CA1 area decreased the number of PV+/PNN+ and total PV+ cells in NTg mice (NTg- ChABC-SH) compared to the control injected mice (NTg-PBS-SH). Tg2576 mice exhibit reduced cell numbers compared to NTg-PBS mice. *: p < 0.05, **: p < 0.01, ***: p < 0.001, Two-way ANOVA with Tukey’s post hoc test. **(C)** Microphotographs illustrating PV (green) and PNN (WFA, yellow) in the CA1 area of NTg mice under SH condition, highlighting PNN degradation 10 days after ChABC injection. **(D)** In NTg-EE mice, ChABC injection did not affect the number of PV+/PNN+ or total PV+ cell in the CA1 area. In Tg2576 mice, EE restored PV+/PNN+ or total PV+ cell counts following PBS injection, but ChABC injection prior to EE prevented this restoration. *: p < 0.05, **: p < 0.01, ***: p < 0.001, Two-way ANOVA with Tukey’s post hoc test. **(E)** PV+/PNN+ and total PV+ cell counts in the CA1 area of Tg2576 mice are compared across different housing and injection conditions. Cell counts are expressed as a percentage of the mean observed in Tg2576 mice housed in standard conditions and injected with PBS solution (PBS-SH). PV+/PNN+ and PV+ cell numbers were restored by EE in PBS-injected Tg2576 mice, but this rescue was prevented by ChABC injection prior to EE. ***: p < 0.001, One-way ANOVA with Dunnett’s post hoc test. **(B,D,E)** Bar graphs show means and SD.

However, there were no differences in PV+/PNN+ and PV+ cell numbers in both CA1 and CA2 areas in NTg mice after 10 days of enriched environment (EE), regardless of PBS or ChABC treatment (**Fig.1D&S1C**). One can assume that the PNNs degraded by ChABC were reformed following 10 days of EE, and therefore no differences between NTg-PBS-EE and NTg-ChABC-EE animals were observed. In Tg2576 mice injected with the control solution, 10 days of EE induced an increased presence of PV+/PNN+ and PV+ cells in CA1 compared to the Tg2576- ChABC-EE mice (**Fig.1D**, two-way ANOVA, PV+/PNN+, interaction: F(1,15) = 9.448; p = 0.0077, Tukey’s post-hoc test: Tg2576-PBS-EE *vs* Tg2576-ChABC-EE, p = 0.0017; two-way ANOVA, interaction: PV+, F(1,15) = 15.33; p = 0.0014, Tukey’s post-hoc test: Tg2576-PBS-EE *vs* Tg2576-ChABC-EE, p = 0.0020). Specifically, after 10 days of EE, Tg2576-PBS-EE mice showed an increase of 227.8% [117.6 to 337.9] of detectable PV+/PNN+ and 227.8% [143.4 to 327.7] in PV+ cells compared to Tg2576-PBS mice that stayed in SH. Thus, 10 days of EE induced a significant increase of PV and PNN presence in the CA1 of Tg2576 mice. Moreover, ChABC injection in CA1 effectively prevented the remodeling of PV and PNN in Tg2576 mice during the 10-day EE period (**Fig.1D&E**).

To evaluate whether these EE-induced changes in PV and PNN numbers lasted at least a couple of weeks, we quantified the number of PV+/PNN+ cells in the CA1 (**Fig.2**) and CA2 (**Fig.S2**) areas of the hippocampus of mice 20 days after the end of the 10-day EE period. These quantifications were therefore carried out 30 days after the injection of PBS or ChABC (**Fig.2A**). For individuals who remained in SH for 30 days, we still observe the expected difference in the number of PNNs around PV interneurons (611.50 [460.30 to 762.80] PV+/PNN+ cells) between NTg and Tg2576 mice, independent of ChABC injection into CA1 area (**Fig.2B&C**, two-way ANOVA, genotype: F(1,48) = 66.09; p < 0.0001). This was accompanied by a decrease in the number of quantifiable PV+ cells (705.00 [510.90 to 899.10] PV+ cells) (**Fig.2B&D**, two-way ANOVA, genotype: F(1,48) = 53.34; p < 0.0001). Moreover, if after 10 days the injection of ChABC in the CA1 area of NTg-SH mice causes a decrease in the number of PV+ and PV+/PNN+ cells, this effect was abolished 30 days post-injection (**Fig.2C&D**). In agreement with the literature ^33,34^ and our unpublished observations, this suggests that 30 days after ChABC injection, the PNNs around the PV cells of NTg mice have started to regrow. This also suggests that the activity of PV cells is not affected in the long term by the ChABC injection.

**Figure 2:**
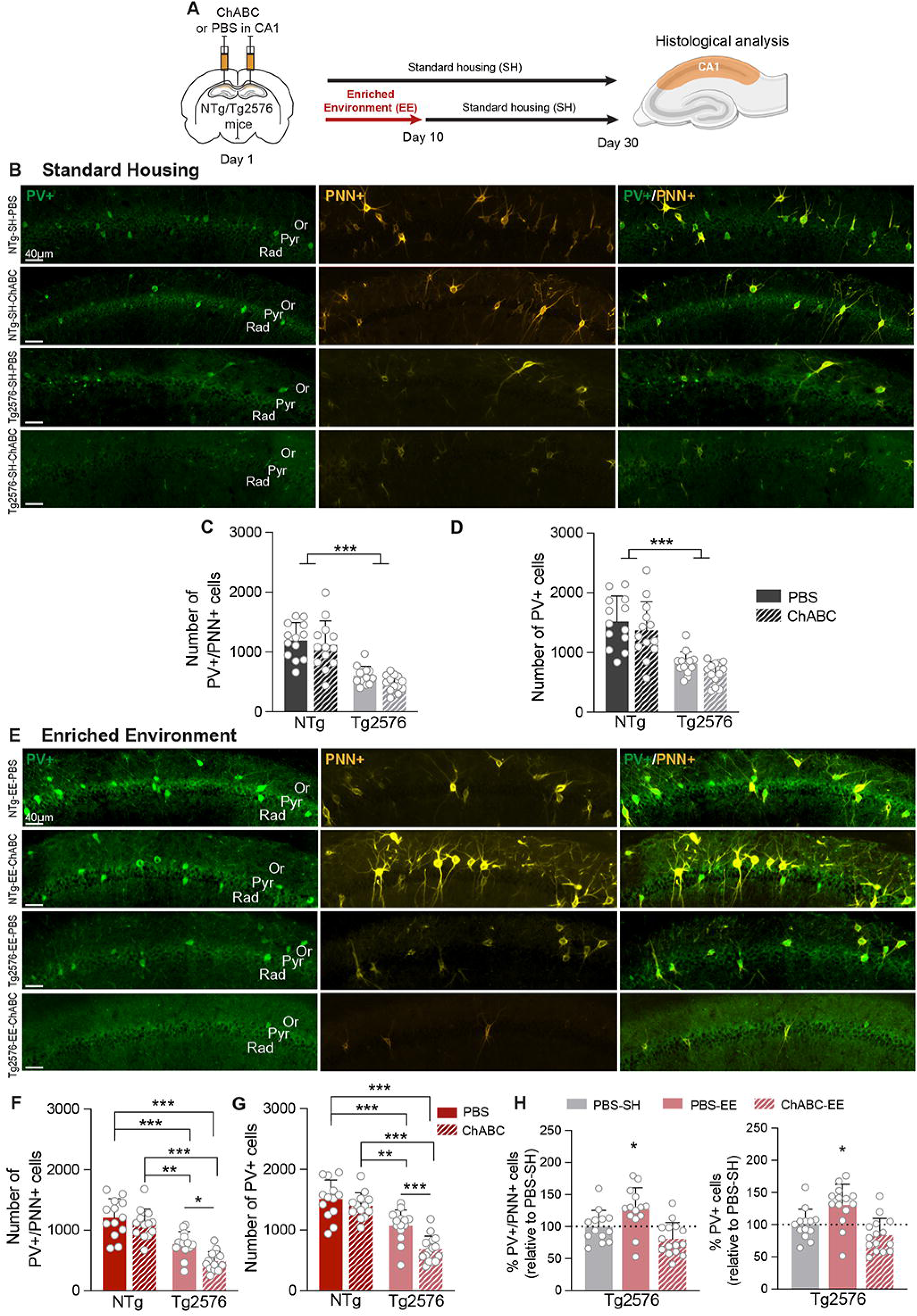
ChABC Injection Into the CA1 Area in Tg2576 Mice Durably Prevents the Restoration of PV+/PNN+ Cells Induced by EE. **(A)** Experimental timeline: ChABC or PBS was injected into the CA1 area of NTg and Tg2576 mice. Mice were either maintained in SH or placed for 10 days in EE before returning to SH. Thirty days after the injections, mice were sacrificed for anatomical quantification**. (B)** Microphotographs illustrating PV+ (green) and PNN+ (WFA staining, yellow) in the CA1 area of NTg and Tg2576 mice that were maintained in SH conditions and injected with either PBS or ChABC. Or: Oriens; Pyr: Pyramidale; Rad: Radiatum. **(C-D)** The number of PV+/PNN+ (C) and total PV+ cells (D) is reduced in Tg2576 mice compared to NTg in SH condition, regardless of the treatment. ***: p < 0.001, Two-way ANOVA, followed by Tukey’s post hoc test. Bar graphs show means and SD. **(E)** Microphotographs illustrating PV+ (green) and PNN+ (WFA staining, yellow) in the CA1 area of NTg and Tg2576 mice, injected with PBS or ChABC, that experienced 10 days of EE following injections. **(F-G)** In mice exposed to 10 days of EE, an increase is observed in PV+/PNN+ (F) and total PV+ (G) cell counts in Tg2576 mice with PBS injection, but not with ChABC injection in the CA1 area. Despite this increase, cell numbers in Tg2576 mice remain below those of NTg mice, regardless of the treatment. **(H)** PV+/PNN+ and total PV+ cell numbers in the CA1 area of Tg2576 mice are compared across housing and injection conditions. A 10-day EE increased PV+/PNN+ and PV+ cell numbers, and this effect lasts for at least 20 days. However, ChABC injection into the CA1 area prior to EE prevented this restoration. p < 0.05, One-way ANOVA with Dunnett’s post hoc test.

Interestingly, 20 days after the end of the EE period, there was a reduction in the number of PNNs around the PV cells of Tg2576 mice compared to NTg-PBS-EE mice, whether they were injected with PBS (436.20 [196.90 to 675.50] cells) or ChABC (715.60 [480.10 to 951.00] cells) (**Fig.2E&F**, two-way ANOVA, genotype: F(1,51) = 68.92; p < 0.0001; injection: F(1,51) = 9.12; p = 0.0039; interaction: F(1,51) = 1.962; p = 0.1673; Tukey’s post-hoc test: NTg-PBS-EE *vs* Tg2576-PBS- EE, p < 0.0001 and NTg-PBS-EE *vs* Tg2576-ChABC-EE, p < 0.0001). This was accompanied by a decrease in the number of detectable PV+ interneurons for the Tg2576-PBS-EE mice (438.40 [178.40 to 698.40] cells) and Tg2576-ChABC-EE mice (824.70 [568.50 to 1080.00] cells) (**Fig.2E&G**, two-way ANOVA, genotype: F(1,51) = 70.54; p < 0.0001; injection: F(1,51) = 13.00; p = 0.0007; interaction: F(1,51) = 4.060; p = 0.0492; Tukey’s post-hoc test: NTg-PBS-EE *vs* Tg2576-PBS- EE, p = 0.0002 and NTg-PBS-EE *vs* Tg2576-ChABC-EE, p < 0.0001). Finally, compared to Tg2576-PBS-SH, we observed an increase of 26.67% [1.52 to 51.81] of PNNs surrounding PV+ interneurons and 30.77% [6.39 to 55.15] of PV+ cells in Tg2576-PBS-EE mice (**Fig.2H**, one-way ANOVA, PV+/PNN+, F(2,39) = 9.451; p = 0.0005; Dunnett’s post-hoc test: Tg2576-PBS-SH *vs* Tg2576-PBS-EE, p = 0.0364; one-way ANOVA, PV+, F(2,39) = 10.80; p = 0.0002; Dunnett’s post-hoc test: Tg2576-PBS-SH *vs* Tg2576-PBS-EE, p = 0.0118), indicating that the 10-day EE period effectively and durably enhanced the presence of PV+/PNN+ and PV+ cells in the area CA1 of Tg2576 mice. Consistent with our previous observations, this increase was not observed in Tg2576-ChABC-EE mice.

Importantly, ChABC injection in area CA1 did not restrain PV/PNN remodeling in the adjacent hippocampal area CA2 of Tg2576 mice following EE (**Fig.S2**).

### ChABC injection into CA1 prevents spatial memory improvement following EE in Tg2576 mice

The same housing protocol was applied to another batch of NTg and Tg2576 mice, 30 days before being tested for their spatial (depending on CA1) and social (depending on CA2) memory (**Fig.3A&4A**). We hypothesize that if PV/PNN remodeling is the substrate for behavioral improvement in Tg2576 mice after EE, then preventing this remodeling with ChABC injection in the CA1 area should specifically preclude any improvement of spatial memory.

**Figure 3:**
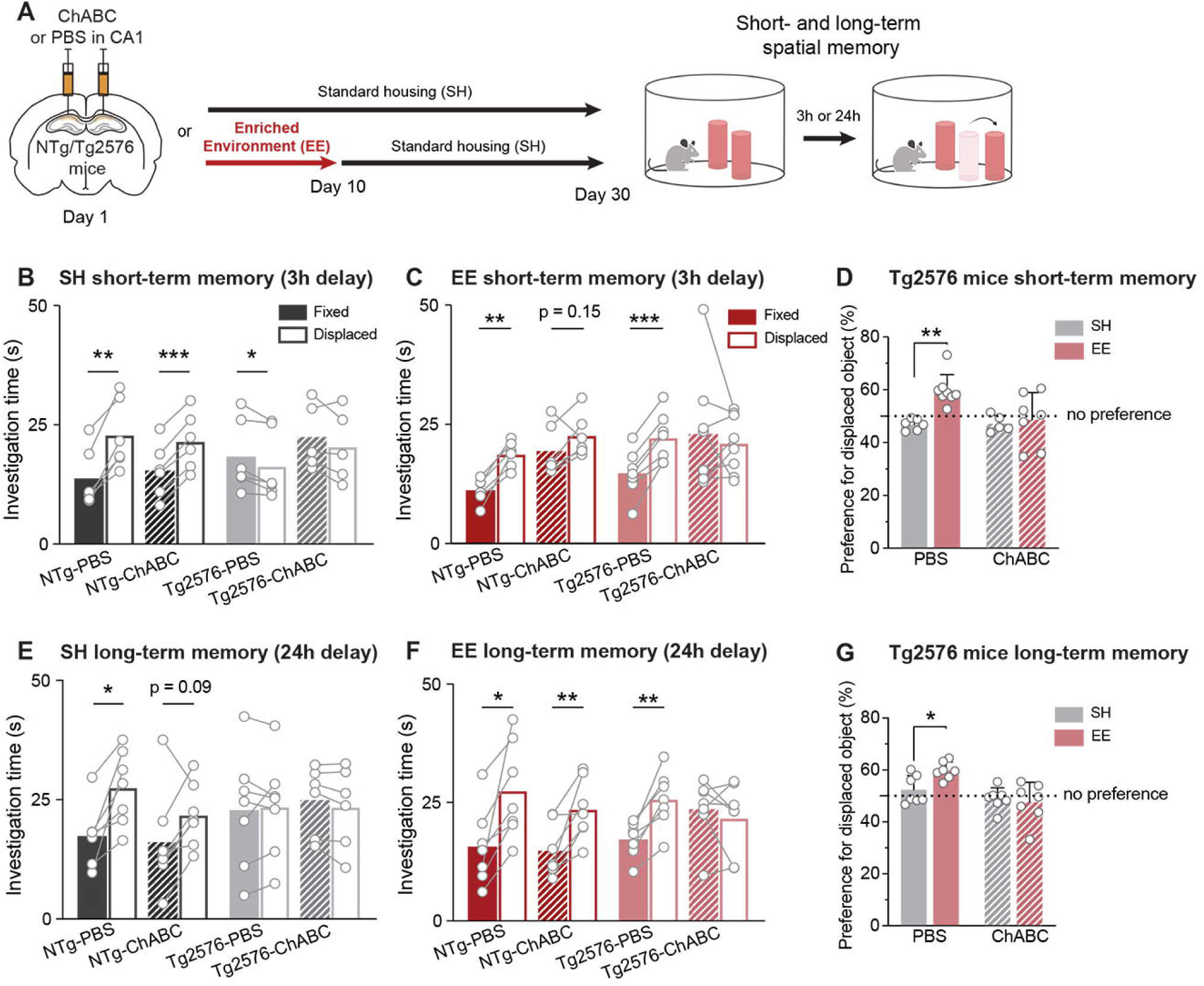
ChABC Injection Into CA1 Prevents Spatial Memory Improvement Following EE in Tg2576 Mice. **(A)** Experimental timeline: ChABC or PBS was injected into the CA1 area of NTg and Tg2576 mice. Mice were either maintained in SH or placed for 10 days in EE before returning to SH. Thirty days after the injections, mice were tested for short- (B-C) and long-term (D-E) spatial memory using the object location test. **(B)** NTg mice maintained in SH spend more time exploring the displaced object, regardless of PBS or ChABC injection, indicating intact short-term spatial memory. In contrast, Tg2576 mice show no preference for the displaced object, indicating impaired spatial memory. **(C)** Tg2576 mice injected with PBS and exposed to a 10-day EE exhibit spatial memory, as shown by a preference for the displaced object. This benefit is lost if ChABC was injected into the CA1 area prior to EE. *: p < 0.05, **: p < 0.01, *: p < 0.001, paired t-test. Each pair of dots with a line represents the investigation time for individual mice. **(D)** Percentage of preference for the displaced object in Tg2576 mice under different housing and treatment conditions. *: p < 0.05: p < 0.01; Two-way ANOVA with Dunnett’s post hoc test. Bar graphs show means and SD. The dotted line indicates no object preference. **(E-F)** Long-term memory object location test for SH (E) or EE (F) conditions showed the same results as for shorter retention delay. *: p < 0.05, **: p < 0.01, paired t-test. **(G)** Percentage of preference for the displaced object in Tg2576 mice across different housing and treatment conditions shows similar trends to the short-term memory test. *: p < 0.05, Two-way ANOVA with Dunnett’s post hoc test.

In the first cohort of mice, short-term spatial memory (STM) was assessed. NTg mice that stayed in SH and were injected with PBS or ChABC were more likely to explore the displaced object (**Fig.3B**, paired t-tests, NTg-PBS-SH fixed 13.78s ± 6.17 *vs* displaced 22.85s ± 7.27 object; t = 6.4145; p = 0.0014 and NTg-ChABC-SH fixed 15.45 s ± 5.68 *vs* displaced 21.51 ± 5.91 object; t = 16.9023; p < 0.0001). This suggests that disrupting PNNs in CA1 30 days before the novel object location test does not impair short-term spatial memory in NTg mice.

In contrast, Tg2576-SH mice had a similar exploration time between the two objects, independent of the injection, *i.e.*, PBS (**Fig.3B**, paired t-tests, Tg2576-PBS- SH fixed 18.35s ± 7.53 *vs* displaced 16.31s ± 7.25 object t = 3.7026; p = 0.0140) or ChABC (Tg2576-ChABC-SH fixed 22.46s ± 6.97 *vs* displaced 20.40s ± 7.50 object; t = 1.7043; p = 0.1635), confirming that 6-month-old Tg2576 mice are impaired in spatial memory, even with a short retention delay.

Interestingly, Tg2576 mice that underwent a 10-day stay in EE after receiving a control solution injection in CA1 showed an improvement in short-term spatial memory (**Fig.3C**, paired t-tests, Tg2576-PBS-EE fixed 14.75s ± 4.94 *vs* displaced 22.19s ± 5.12 object; t = 8.9154; p < 0.0001). Specifically, there was a 13.98% [5.07 to 22.88] increase in preference for the displaced object compared to Tg2576 mice that stayed in SH (**Fig.3D**, two-way ANOVA, interaction: F(1,22) = 5.585; p = 0.0274; Dunnett’s post-hoc test: Tg2576-PBS-SH *vs* Tg2576-PBS-EE, p = 0.0019).

This indicates that 10 days of EE are sufficient to induce changes that last at least 20 days and sustain short-term spatial memory improvement.

However, Tg2576 mice that were injected with ChABC in CA1 before EE did not show any improvement in short-term spatial memory (**Fig.3C&D**, paired t-tests, Tg2576-EE-ChABC fixed 23.00s ± 12.18 *vs* displaced 21.03s ± 6.11 object t = 0.5879; p = 0.5751), suggesting that preventing PV/PNN remodeling during SEE through PNN degradation precludes EE-dependent memory improvement.

To examine whether EE also has a beneficial impact on spatial memory consolidation in Tg2576 mice, another set of mice was tested with a 24-hour delay between acquisition and test (long-term memory, LTM). In SH condition, NTg mice injected with PBS spent more time exploring the displaced object than the fixed one (**Fig.3E**, paired t-tests, NTg-PBS-SH fixed 17.40s ± 6.40 *vs* displaced 27.50s ± 7.77 object; t = 3.7542; p = 0.0095). In NTg mice injected with ChABC, recall seemed slightly disturbed, mainly due to one individual spending more time interacting with the object at the familiar location (NTg-ChABC-SH fixed 16.12s ± 10.74 *vs* displaced 21.77s ± 6.64 object; t = 2.0021; p = 0.0922).

Consistent with our observations at a shorter memory delay, Tg2576 mice injected with PBS and placed in EE spent more time exploring the displaced object (**Fig.3F**, paired t-tests, Tg2576-PBS-EE fixed 17.22 s ± 3.74 *vs* displaced 25.70 s ± 6.02 object; t = 5.7917; p = 0.0012), with a 7.42% [0.18 to 14. 66] increase in preference over Tg2576-SH-PBS mice (**Fig.3G**, two-way ANOVA, interaction: F(1,24) = 4.515; p = 0.0441; Dunnett’s post-hoc test: Tg2576-PBS-SH *vs* Tg2576-PBS-EE, p = 0.0437). This result confirms that a transient stay in EE improves the spatial memory of Tg2576 mice, even with a 24-hour recall.

However, as observed for short-term memory, the 10-day stay in EE did not restore long-term spatial memory in Tg2576 mice if they received an injection of ChABC in CA1 area before EE (**Fig.3F**, paired t-tests, Tg2576-ChABC-EE fixed 23.60s ± 6.66 *vs* displaced 21.69s ± 7.60; t = 0.8678; p = 0.4189). This experiment demonstrates that the beneficial effect of EE on spatial memory in Tg2576 mice is maintained when CA1-related memory requires consolidation. These results indicate that blocking the remodeling of PV interneurons and their PNNs in the CA1 area during EE prevents the EE-dependent cognitive benefits on the spatial memory of Tg2576 mice.

### ChABC injection into CA1 area still allows EE-induced improvement of social memory in Tg2576 mice

The fact that ChABC injection into CA1 of Tg2576 mice prevents EE-induced beneficial effects on spatial memory suggests that this CA1-dependent memory improvement is sustained by the increased number of detectable PV+ cells following EE. Therefore, in these same mice, the ChABC-induced perturbation of PV/PNN in CA1 should not alter social memory capacity, which is dependent on the integrity of PV function in area CA2 ^24,29,35^.

We first examined whether ChABC injection into CA1 impacted social memory performance in both NTg and Tg2576 mice housed in standard conditions. Using the 5-trial test assessing short-term social memory, we observed that 6-month-old NTg and Tg2576 PBS-injected mice behaved differently during the four exposures to the same mouse (**Fig.4A**). NTg-SH-PBS mice showed a significant decrease of 40.69% [18.86 to 62.51] in interaction time with the exposed mouse by the second trial (**Fig.4B**, one-way ANOVA, NTg-SH-PBS, trial: F(3,32) = 19.93; p < 0.0001; Dunnett’s post-hoc test: trial 1 *vs* trial 2, p = 0.0007). Social learning was slower in Tg2576-SH-PBS mice (**Fig.S3B**), where a significant decrease of 45.01% [19.34 to 70.68] in interaction time was observed only at the third trial (**Fig.4B**, one-way ANOVA, Tg2576-SH-PBS, trial: F(2,21) = 5.447; p = 0.0133; Dunnett’s post-hoc tests: trial 1 *vs* trial 2, p = 0.5928; trial 1 *vs* trial 3, p = 0.0014). Therefore, the social memory ability of Tg2576 mice is already altered at 6 months of age, although this deficit is not as profound as observed in 9-month-old individuals ^29^.

**Figure 4.**
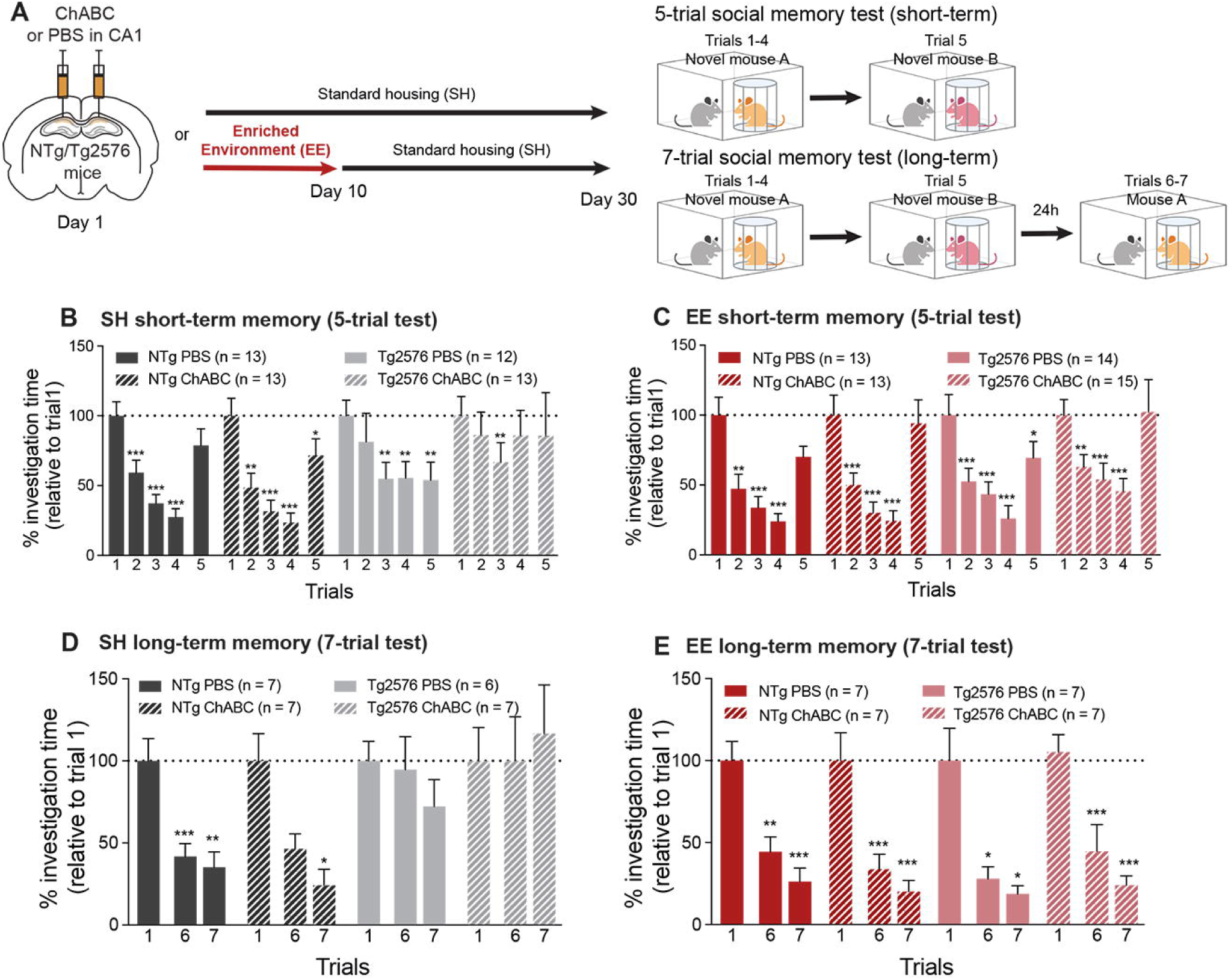
PNN Disruption in Area CA1 Does Not Impair EE-Induced Improvement of Social Memory Capacity in Tg2576 Mice. (A) Experimental timeline: Animals were injected with PBS or ChABC into the CA1 area, then exposed to an EE for 10 days followed by SH for 20 days, or remained in SH for 30 days. Afterward, mice were tested for short-term (5-trial) and long-term (7-trial) social memory. **(B-C)** Percentage of interaction time relative to trial 1 in SH (B) and EE (C) conditions for the 5-trial test. Tg2576 mice show impaired social memory when maintained in SH condition. Ten days of EE after either PBS or ChABC injection into the CA1 area restored social memory of Tg2576 mice, which lasts for at least 20 days after the end of EE. *: p < 0.05, **: p < 0.01, ***: p < 0.001, one-way ANOVA with Dunnett’s post hoc test. **(D-E)** Long-term social memory, assessed with the 7-trial test, was impaired in Tg2576 mice in SH conditions (D). Exposure to EE (E) rescued long-term social memory in Tg2576 mice, regardless of treatment. *: p < 0.05, **: p < 0.01, ***: p < 0.001, one-way ANOVA with Dunnett’s post hoc test. Bars represent means, and error bars indicate SEM. The dotted line indicates 100%, reflecting the interaction time during trial 1.

Interestingly, ChABC injection in area CA1 did not modify the differences in social memory ability observed in standard-housed NTg and Tg2576 mice (**Fig.4B**, one-way ANOVA, NTg-SH-ChABC, trial: F(2,21) = 22.22; p < 0.0001; post-hoc Dunnett’s test: trial 1 *vs* trial 2, p = 0.0030, 51.38% [18.62 to 84.13]; one-way ANOVA, Tg2576-ChABC-SH, trial: F(2,20) = 0.738; p = 0.4791; Dunnett’s post-hoc test : trial 1 *vs* trial 2, p = 0.7399). Remarkably, after the EE period, Tg2576 mice showed a reduction in investigation time from trial 2, demonstrating that EE restored social memory in Tg2576 mice, independently of ChABC injection in the CA1 area (**Fig.S3C&4C**).

Furthermore, another set of mice was tested for long-term social memory, *i.e.*, re-exposed the following day to the mouse presented during the first 4 trials. We observed that the NTg mice displayed a reduced interaction time with the now familiar mouse during trials 6 and 7, independent of ChABC injection and housing conditions (**Fig.4D&E**, one-way ANOVA, NTg-PBS-SH, trial: F(2,9) = 23.24; p = 0.0005; Dunnett’s post-hoc test: trial 1 *vs* trial 6, p = 0.0007; one-way ANOVA, NTg- ChABC-SH, trial: F(1,5) = 10.89; p = 0.0162; Dunnett’s post-hoc test: trial 1 *vs* trial 6, p = 0.1146; one-way ANOVA, NTg-PBS-EE, trial: F(2,9) = 31.33; p < 0.0001; Dunnett’s post-hoc test: trial 1 *vs* trial 6, p = 0.0052; one-way ANOVA, NTg-ChABC- EE, trial: F(1,7) = 45.93; p = 0.0002; Dunnett’s post-hoc test: trial 1 *vs* trial 6, p = 0.0005). As expected, Tg2576-PBS-SH and Tg2576-ChABC-SH mice showed similar interaction times during the 6th and the 1st trial (**Fig.4D**, one-way ANOVA, Tg2576-PBS-SH, trial: F(2,9) = 1.188; p = 0.3420; Tg2576-ChABC-SH, trial: F(2,9) = 0.1439; p = 0.8459), indicating their inability to memorize in the long term a mouse presented to them the day before.

Remarkably, the impairment in the social memory test of 6-month-old Tg2576 mice was completely abolished by a 10-day stay in EE, regardless of whether they had received an injection of ChABC in the CA1 area (**Fig.4E**, one-way ANOVA, Tg2576- PBS-EE, trial: F(1,7) = 12.96; p = 0.0063; Dunnett’s post hoc test: trial 1 *vs* trial 6, p = 0.0264; one-way ANOVA, Tg2576-ChABC-EE, trial: F(2,11) = 34.82; p < 0.0001; Dunnett’s post hoc test: trial 1 *vs* trial 6, p = 0.0008). This indicates that EE-induced environmental stimulations had a beneficial effect on CA2-related social memory capacity of Tg2576 mice, and that limiting PV/PNN remodeling in CA1 during EE does not prevent this beneficial effect. Therefore, EE has a beneficial impact on short- and long-term social memory of 6-month-old Tg2576 mice, independent of PBS or ChABC injection into CA1.

Altogether, these data suggest that preventing PV/PNN remodeling in CA1 during EE with ChABC injection abolishes the spatial memory improvement due to EE in Tg2576 mice, but not their CA2-related social memory enhancement.

### NRG1 injection into CA1 restores spatial memory in Tg2576 mice

Our results strongly suggest that the remodeling of the PV/PNN network due to EE is necessary to the improvement of hippocampal-dependent memory in Tg2576 mice. Another way to induce the formation of PNNs around the PV cells is to locally inject the neurotrophic factor neuregulin-1 (NRG1) ^29,36^. Here, we aimed to use NRG1 to promote the maturation of PNN around PV+ cells locally in the CA1 area and observe whether it was sufficient to restore spatial memory capacity in Tg2576 mice (**Fig.5A**).

**Figure 5:**
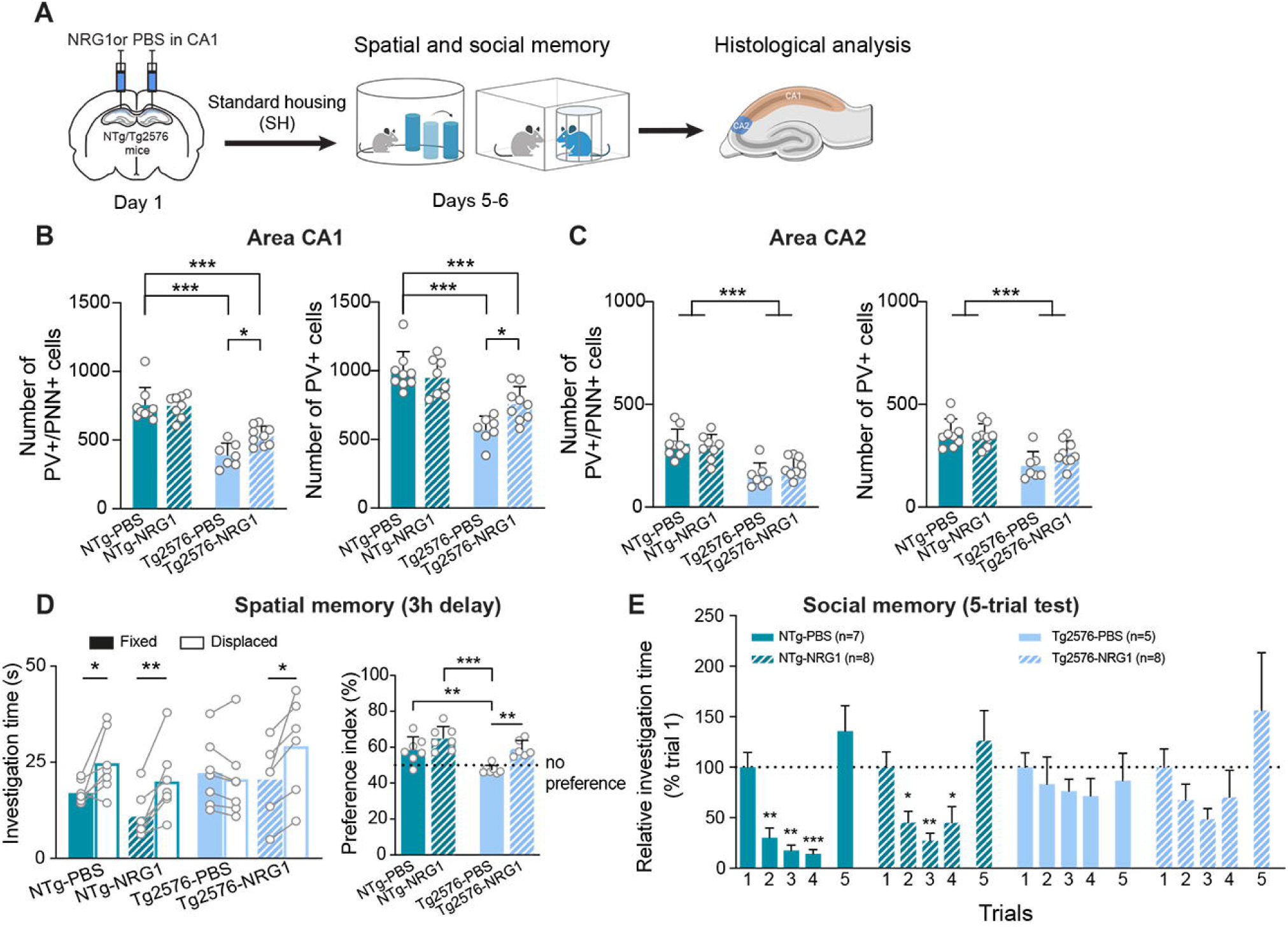
Local NRG1 Injection in CA1 Area Restores PV+/PNN+ Cells in Tg2576 Mice. **(A)** Experimental timeline: Mice were injected with either PBS or NRG1 in the CA1 area. Five days post-injection, spatial and social memory were assessed. Following memory tests, brains were collected to quantify PV+/PNN+ and PV+ cells in the CA1 and CA2. **(B)** Quantification of PV+/PNN+ and PV+ cells in the CA1 area. NRG1 injection significantly increased the number of PV+ cells and restored the presence of PNN around PV cells in Tg2576 mice, though not to levels observed in NTg mice. *: p < 0.05, ***: p < 0.001, Two-way ANOVA with Tukey’s post hoc test. Bar graphs show means and SD. **(C)** In the CA2 area, NRG1 injection did not restore the number of PV+/PNN+ or PV+ cells, indicating that the restorative effects of NRG1 are specific to the CA1 area. ***: p < 0.001, Two-way ANOVA with Tukey’s post hoc test. Bar graphs show means and SD. **(D)** Spatial memory was assessed using the object location test. Tg2576 mice exhibited impaired spatial memory, and NRG1 injection into the CA1 area significantly increased the time spent exploring the displaced object (left). *: p < 0.05, **: p < 0.01, Paired t-test. The preference index (right) further showed that NRG1 injection in Tg2576 mice restored spatial memory, compared to those injected with PBS. **: p < 0.01; ***: p < 0.001, Two-way ANOVA with Dunnett’s post hoc test. Bar graphs show means and SD. The dotted line indicates no object preference. **(E)** Percentage of investigation time in the 5-trial test relative to trial 1. Tg2576 mice exhibit impaired social memory, which was not improved by NRG1 injection into CA1. *: p < 0.05, **: p < 0.01, ***: p < 0.001, one-way ANOVA with Dunnett’s post hoc test. Bars represent means and SEM. The dotted line indicates 100%, reflecting the interaction time during trial 1.

Five days after the bilateral injection of NRG1 into the CA1 area of 9-month-old mice, we performed a quantification of PV+/PNN+ cells in the CA1 and CA2 areas **(Fig.5A)**. In Tg2576-NRG1 mice, we observed an increase of 140.5 [8.331 to 272.40] PV+/PNN+ cells compared to Tg2576-PBS mice (**Fig.5B**, two-way ANOVA, PV+/PNN+, interaction: F(1,29) = 4.624; p=0.0400; Tukey post-hoc test: Tg2576- PBS *vs* Tg2576-NRG1, p = 0.0339). This increase in PV+/PNN+ cells in Tg2576- NRG1 mice was accompanied by an increase of 187.32 [10.39 to 364.21] detectable PV+ cells in CA1 compared to Tg2576 mice injected with PBS (**Fig.5B**, two-way ANOVA, PV+, interaction: F(1,29) = 6.751; p = 0.0146; Tukey’s post-hoc test: Tg2576-PBS *vs* Tg2576-NRG1, p = 0.0349). Thus, the histological analysis demonstrates that a single injection of NRG1 is sufficient to increase the number of detectable PV+ and PV+/PNN+ neurons in the CA1 area of Tg2576 mice. In contrast, in area CA2, we observed only the expected difference between NTg and Tg2576 mice for PV+/PNN+ cells (**Fig.5C**, two-way ANOVA, genotype: F(1,29) = 34.27; p < 0.0001) and for PV+ cells (**Fig.5C**, two-way ANOVA, genotype: F(1,29) = 29.64; p < 0.0001). The injection of NRG1 into CA1 had no effect on the number of detectable PV+/PNN+ and PV+ cells in CA2, regardless of the genotype of the mice.

Short-term spatial memory of NTg and Tg2576 mice was assessed usingthe object location test. NTg mice were more likely to explore the displaced object, regardless of the injections (**Fig.5D**, paired t-tests, NTg-PBS: fixed 16.99s ± 2.93 *vs* displaced 24.77s ± 8.10 object; t = 2.698; p = 0.0357; NTg-NRG1: fixed 10.86s ± 5.81 *vs* displaced 20.03s ± 9.234 object; t = 4.551; p = 0.0039). This suggests that NRG1 injection in CA1 does not influence the spatial memory of NTg mice. In contrast, Tg2576 mice injected with PBS showed similar exploration times between the two objects (**Fig.5D**, paired t-test, Tg2576-PBS: fixed 22.17s ± 8.85 *vs* displaced 20.60s ± 10.27 object; t = 1.571; p = 0.1672). Consistent with what we observed after remodeling of the PV/PNN network following a 10-day EE, Tg2576 mice injected with NRG1 were able to discriminate the displaced from the fixed object (**Fig.5D**, paired t-test, Tg2576-NRG1: fixed 20.53s ± 9.92 *vs* displaced 29.20s ± 13.04 object; t = 3.371; p = 0.0199), with an increase of 11.57% [2.912 to 20.22] compared to the Tg2576-PBS group. (**Fig.5D**, one-way ANOVA, F(3,23) = 11.56; p < 0.0001; Tukey’s post-hoc test Tg2576-PBS *vs* Tg2576-NRG1, p = 0.0061). Hence, inducing PV/PNN remodeling in CA1 by NRG1 injection restores the spatial memory capacities of Tg2576 mice.

When subjected to the 5-trial social memory test, which relies on CA2 area, NTg mice injected with PBS or NRG1 showed a significant decrease in interaction time of 69.82 % [25.81 à 113.8] and 54.78 % [22.31 to 87.26], respectively, as early as the second trial (**Fig.5E**, one-way ANOVA, NTg-PBS, trial: F(2,10) = 15.38; p = 0.0017; Dunnett’s post-hoc test : trial 1 *vs* trial 2, p = 0.0066; one-way ANOVA, NTg-NRG1, trial: F(2,10) = 8.305; p = 0.0144; Dunnett’s post-hoc test: trial 1 *vs* trial 2, p = 0.0039). In Tg2576-PBS mice, there was no difference in interaction time between the trials compared to the first one (**Fig.5E**, one-way ANOVA, trial: F3,11 = 0.6395; p = 0.5931), and NRG1 injection in CA1 did not affect this CA2-related memory capacity in 6-month-old Tg2576 (**Fig.5E**, one-way ANOVA, trial: F(2,12) = 1.929; p = 0.1911). Altogether, these results indicate that the remodeling of the PV/PNN network with NRG1 injection in CA1 improves memory related to this specific area, but not other hippocampal-dependent memory.

## Discussion

Our study demonstrates that a 10-day period of exposure to an enriched environment effectively restores both spatial and social memory in Tg2576 mice, along with an increase in the population of PV+ and PV+/PNN+ cells within the hippocampus. However, the injection of ChABC into the CA1 area disrupts this restoration by preventing the remodeling of PV interneurons and their PNNs, thereby blocking the improvement in spatial memory. Conversely, injecting NRG1 into the CA1 area of Tg2576 mice induces the remodeling of PV interneurons and their PNNs, specifically restoring spatial memory. These findings establish a direct link between the remodeling of PV cells in the hippocampus due to environmental stimulation and the resulting cognitive benefits in an AD mouse model.

To inhibit the remodeling of PV/PNN neurons, we employed the enzyme ChABC, which degrades the PNNs around PV cells when injected into specific brain regions. ChABC, a bacterial enzyme from *Proteus vulgaris* ^37^, digests the chondroitin sulfate (CS) chain into its disaccharide units, removing the glycosaminoglycan (GAG) chain from the core protein. Although there is uncertainty about the exact effects of ChABC on PNNs and surrounding neurons, as well as its specificity for the CS of PNNs, this enzyme has been instrumental in advancing our understanding of plasticity mechanisms in the nervous system ^23,38^. In our study, ChABC injection significantly reduced the number of detectable PV+/PNN+ cells in the CA1 area of NTg mice for at least 10 days, consistent with the duration of EE.

Additionally, we show that a 10-day EE improves hippocampal-dependent memory and triggers the recovery of PV and PNN levels in the hippocampus of Tg2576 mice. Notably, a 10-day EE is as effective as the 10-week EE protocol used in our previous studies ^27,30^. It is important to note that EE duration varies widely across studies on nervous system disorders, often leading to varied outcomes ^31,32^, and questioning the need for standardized EE protocols ^39^. Consistent with our observations, previous research has shown that 10 days of EE are associated with a change in configuration of hippocampal PV interneurons in adult wild-type mice, shifting toward a configuration that is conducive to new experiences and learning. This corresponds to reduction in PV expression due to changes in the number and nature of synaptic connections onto PV neurons ^40^, possibly facilitated by the reduction of PNNs around their soma and perisomatic terminals ^19,27^. As a result, EE modifies PV interneuron activity in the hippocampus, leading to disinhibition of pyramidal neurons and facilitating new learnings ^40^. In contrast, in AD models where the presence and activity of PV/PNNs are significantly altered in control conditions ^27^, restoring the activity of these neurons improves their functions ^16,29^. If EE appears to have opposite effects in healthy mice and pathological models, it is possible that the variety of experiences it provides optimally modulates PV/PNN network into a state that enables cognitive benefits, regardless of their baseline state.

In the context of AD models, we propose that the experiences provided by an EE rapidly enhance the activity of PV neurons. This may be associated with a structural and spatial reorganization of the extracellular matrix, leading to the maturation of PNNs around PV neurons ^41^. The reorganization of specific PNN components could have distinct and beneficial effects on cognitive functions. For instance, correlations have been reported between the presence of brevican and improvements in spatial working and short-term memory ^19,42^. However, it remains to be determined whether it is the PNN components or the activity of PV cells that primarily drive this remodeling, influencing both the structural organization of the PNN and PV cell activity.

PNN degradation and plasticity are regulated by various proteases, including matrix metalloproteinases (MMPs) and ADAMTS ^43–45^. These proteases are elevated in the brains of AD mouse models ^46^ and patients ^47^. Specifically, MMP9 significantly modulates PNNs in the hippocampus by degrading many of their components ^48,49^. In wild-type mice, MMP-9 levels increase around PV interneurons with sparse PNNs after EE exposure, suggesting its role in reducing PNN presence following EE ^50^. In Tg2576 mice, the baseline state of the network might cause EE to have an opposite effect on MMP9, reducing its activity and thus allowing PNN formation around PV neurons.

We hypothesize that mechanisms involved in PNN maturation during juvenile stages might also play a role in restoring this network in Tg2576 mice. Previous studies have shown that the activity of excitatory neurons triggers the release of NRG1 ^51,52^, which promotes the maturation of PV interneurons and PNNs ^53^ through NRG1/*ErbB4* signaling. By injecting NRG1, we were able to replicate the effects of EE locally, enhancing PV neuron activity and supporting our hypothesis that hippocampus-dependent memory can be specifically restored by stimulating PV neuron activity and promoting PNN maturation in Tg2576 mice ^29^. The strong expression of the ErbB4 receptor strongly in PV cells, which coincides with the presence of PNNs ^24^, further supports this mechanism. Additionally, our study suggests that ChABC injections might reduce ErbB4 receptor levels on PV interneurons.

Our study aimed to establish a causal link between PV cell remodeling in the hippocampus, driven by environmental stimulation, and cognitive benefits in Tg2576 mice. We propose that PV interneurons and their PNNs are central to the cognitive benefits induced by EE, as supported by existing literature.

Conceptually, in humans, EE is analogous to the concept of cognitive reserve, although its underlying substrates are challenging to identify. Cognitive reserve is defined by lifestyle factors that protect individuals against the development of neuropathology and/or the progression of cognitive decline ^54–56^. We suggest that life experiences of humans, or EE for non-human animals, strengthen the activity of PV GABAergic neurons and potentially contribute to the development of cognitive reserve. This concept has been explored through the use of photic and other sensory driving stimulation of cortical PV interneurons to assess cognitive benefits in AD mouse models^57,58^, and more recently, in patients ^59^. These findings open up possibilities for non-invasive treatments, which still merit further investigation.

## Conclusion

This study demonstrates that environmental enrichment induces the remodeling of PV cells and their PNNs, a process essential for cognitive function improvement in Tg2576 mice, a model of AD. Our findings not only shed light on the mechanisms underlying cognitive enhancement in AD models but also open new research avenues focused on the detailed remodeling of PV interneurons. This includes examining both excitatory and inhibitory synaptic connections, as well as the structural organization of PNNs. By deepening our understanding of PV/PNN neuron functionality, this work paves the way for identifying novel therapeutic targets for neurodegenerative diseases like AD. Furthermore, it enriches our understanding of how environmental stimuli are integrated into neural circuits to support cognitive health.

## Methods

### Mouse model

Experiments comparing housing conditions (standard housing, SH, and enriched environment, EE) were conducted on a total of 153 female mice from the Tg2576 mouse line, aged 5 to 6 months. The experiments involving NRG1 were performed on 33 NTg and Tg2576 female mice, aged 8 and 11 months. All mice were derived from our in-house Tg2576 transgenic colony. This line overexpresses a double mutant form of the human APP695 (Lys670-Asn, Met671-Leu [K670N, M671L]), under the control of the hamster prion protein promoter ^60^. Mice were housed in groups of 2 to 5 per standard home cage and maintained on a 12-hour light/dark cycle with unrestricted access to food and water. The choice to use 5-month-old mice was based on previous findings indicating the onset of learning, cognitive, and memory deficits ^61,62^ as well as cellular ^63,64^ and molecular ^65^ alterations during this period.

### Stereotaxic injections

For perineuronal net (PNN) degradation, 5-month-old female NTg and Tg2576 mice (n=153) were anesthetized using vetflurane (2-5%; Virbac, Cat# Vnr137317) and secured in a stereotaxic frame (Kopf). Before injections, lidocaine (0.1mL at 7mg/kg) was applied subcutaneously to the scalp to provide local anesthesia. Bilateral injections were performed in the CA1 area of the dorsal hippocampus (position relative to bregma: -1.2 mm anteroposterior, ± 1.1 mm mediolateral, -1.6 mm dorsoventral, and -1.8 mm anteroposterior, ± 1.8 mm mediolateral, -1.75 mm dorsoventral). A total volume of 100nL for rostral injections and 200nL for caudal injections was administered, using a solution containing ChABC (50 U/ml, Sigma-Aldrich, Cat# C3667, n=78), or a vehicle solution (0.1M phosphate buffer saline, PBS; n=75). To ensure optimal diffusion, a 5-minute delay was observed after each of the four injections per animal. Following the injections, the scalp was sutured, and the mice received an intraperitoneal injection of metacam (0.1mL) for postoperative analgesia. The animals were placed under a heat lamp for one hour to aid in recovery. The same protocol was used for NRG1 injections (6.66nM; n=16 PBS, n=17 NRG1).

### Enriched environment

One day following stereotaxic injections, both Tg2576 and NTg mice, treated with either ChABC or PBS, were randomly assigned to two housing conditions: standard housing (SH) or an enriched environment (EE). The standard housing group remained in their laboratory cages, with 2-5 mice per cage, until reaching 6 months of age. The EE group was transferred to an enriched environment for 10 days, after which they were returned to their standard home cages for 20 days, until they reached 6 months of age. Cognitive assessments were conducted on both groups, followed by euthanasia for further analysis.

The EE consisted of a large box (100 × 50 × 80 cm) with unlimited access to food and water, and containing a variety of objects differing in shape, size, color, material, and texture, but no running wheels were included. The environment also provided social enrichment through the presence of a larger number of mice (11-16 individuals, from 3 to 5 different cages). To promote exploration and novel experiences, the spatial configuration of the EE was modified every two days with the introduction of new objects, as described in ^66^ and ^30^.

### Behavioral experiments

*Object location test*. The object location test assesses the ability of mice to discriminate between familiar and novel object locations within an open-field (40 cm diameter) that includes a visual cue (striped pattern). The test begins a 10-min habituation phase, allowing the mice to acclimate to the arena. During the acquisition phase, two identical objects are placed equidistant from the walls of the open-field (10 cm), and the mice are given 10 min to freely explore. The time spent exploring each object is recorded. An exclusion criterion was set at a total exploration time of less than 15 seconds (only 1 mouse was excluded based on this criterion). The test phase occurs either 3 hours (short-term) or 24 hours (long-term) after the acquisition, and lasts 10 min. In this phase, one of the two objects is displaced to a new location, pseudo-randomly chosen to minimize bias towards any particular position. Objects and the arena were cleaned with 30% ethanol between trials to eliminate olfactory cues.

*Five-trial social memory test*. This test evaluates the ability of the mice to distinguish between familiar and novel conspecifics by measuring interaction time across multiple trials ^29,67^. Initially, each mouse undergoes a 10-min habituation in a large cage (30 × 60 cm) containing an empty cylindrical cage (8 × 16 cm) positioned in the center. A novel, age- and sex-matched mouse is then introduced into the cylindrical cage for four successive 5-min trials with a 10-min inter-trial interval (ITI). During the fifth trial, a new unfamiliar mouse is introduced into the cylindrical cage. For the long-term memory assessment, 24 hours later, the same mouse from trials 1-4 is presented again in trials 6 and 7. The direct interaction time of the tested mouse with the presented mouse is recorded for each trial. Both the chamber and the small cage are cleaned with 30% ethanol between animals, and the small cage is cleaned between trials to prevent olfactory interference.

### Tissue preparation and immunohistochemistry

Following completion of the behavioral tests, mice were sacrificed on the same day. They were deeply anesthetized using Dolethal solution (150 mg/kg) and then perfused transcardially with 0.9% saline solution for 45-60 seconds. The brains were removed and post-fixed in 4% formaldehyde in a 0.1M phosphate buffer (PB) solution for 48 hours at 4°C. Subsequently, brains were submerged in a 30% sucrose solution containing 0.1% sodium azide for a minimum of 1 week. Coronal brain sections (30µm thick) were obtained using a sliding microtome (Leica SM2010R) equipped with a freezing-stage (Physitemp BFS-3MP). Sections ranging from -0.6 to -3.0 mm relative to bregma were extensively rinsed in PBS and blocked for one hour in a solution containing 10% normal donkey serum (NDS) with PBS and 0.25%Triton-X (PBST), and 0.1 sodium azide. Sections were then incubated overnight at room temperature in NDS solution with biotin-conjugated Wisteria floribunda agglutinin (WFA) lectin (1:1000; Sigma, Cat#L1516), rabbit anti-PCP4 (1:500; Santa Cruz, Cat#sc-74816), and goat anti-PV antibodies (1:2500; Swant, Cat#PVG213). After incubation, sections were washed three times in PBS for 20 min each and then incubated for 90 min at room temperature with 1:1000 streptavidin coupled with TRITC (Vector, Cat#SA-5549) for PNN, 1:500 donkey anti-goat Alexa488 (Invitrogen, Cat#A-11055) for PV cells, and 1:500 donkey anti-rabbit A647 (Invitrogen, Cat#A-31573) for PCP4 cells. Sections were mounted using Mowiol solution and coverslipped for imaging and analysis.

### Image analysis

Quantification of PV-immunoreactive (PV+) and PV+/PNN+ cells was performed on a 1-in-10 series of brain sections, spaced at 300 µm apart, spanning the dorsal hippocampus (beginning at -1.00 mm from bregma). For each mouse, the PV+, PNN+, and PV+/PNN+ cells were manually counted using a fluorescence microscope (Leica DM6000 B) at x20 magnification. Double immuno-labeling for PV and PNN was determined when WFA staining completely enveloped the soma of a PV+ cell, even if the staining intensity was low. Adjustments in focal planes were employed by the experimenter to minimize uncertainty.

The surface area of the hippocampal CA1 and CA2 areas, sampled for counting, was measured using the Mercator stereology system (Explora Nova, La Rochelle, France) at x5 magnification. To enhance the accuracy of CA2 area identification, PCP4 labeling was used, which is specific to the pyramidal cells of the CA2 area in the hippocampus ^68^. The total number of PV+, PNN+ and PV+/PNN+ cells was estimated by multiplying the reference volume of the CA1 and CA2 areas by the density of neurons per sectional volume ^27,69^.

### Ethics statement

All experiments were conducted in strict accordance with the policies of the European Union (2010/63/EU), and the guidelines set forth by the French National Committee of Ethics (87/848 and its modification of February 1st, 2013) for the care and use of laboratory animals. The animal facility of the CRCA is fully accredited by the French Direction of Veterinary Services (E 31-555-011, June 17^th^, 2021), and experimental procedures conducted in this study were approved by local ethical committees and the French Ministry for Research (APAFIS#20210- 2019041014597476 v4). Every effort was made to improve animal welfare and minimize suffering.

### Statistics

Data analysis and graph generation were performed using GraphPad Prism 8 (GraphPad Software Inc., La Jolla, CA, USA). Null hypotheses were rejected at the 0.05 significance level. Group differences are reported with effect sizes corresponding 95% confidence intervals [95% CI]. Results are presented with mean ± standard deviation.

## Acknowledgments

The authors thank Sébastien Gauzin and Maud Combe for technical help. Mice were housed in the ABC Facility of ANEXPLO, Toulouse. The authors greatly acknowledge the Mouse Behavioral Core (MBC) of the Center of Integrative Biology. This work was supported by the Centre National de la Recherche Scientifique (CNRS), the University of Toulouse, the Association France Alzheimer (AAP SM 2018 #1823), the Fondation Vaincre Alzheimer (Grant #61144). G.B. received a PhD fellowship from Toulouse University as part of the “Science of aging” program.

## Author contributions

G.B. and L.V. conceived the project and designed the experiments. G.B., F.J.T., T.D., C.L. and L.V., performed the experiments and analyzed the data. All authors discussed the results, and G.B., F.J.T. and L.V. wrote the manuscript.

## Supplementary Figure Legends

**Figure S1:**
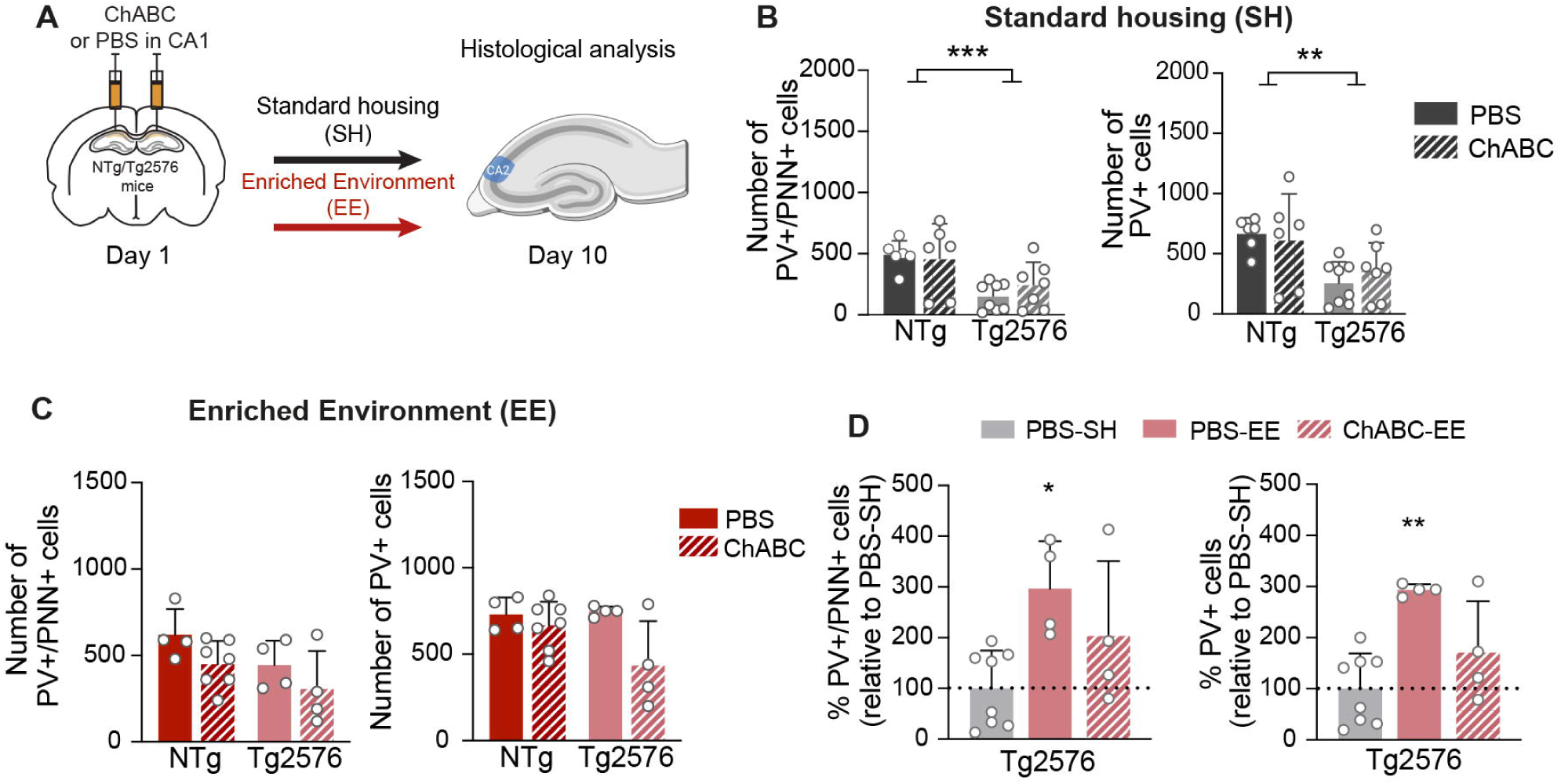
Ten days of EE are sufficient to rescue PV+/PNN+ cells number in area CA2. **(A)** Experimental timeline: ChABC or PBS was injected into CA1 area of NTg and Tg2576 mice, then housed in either SH or EE conditions. Mice were sacrificed 10 days later for anatomical quantifications in the CA2 area. **(B)** ChABC injection into the CA1 area did not affect PV+ or PV+/PNN+ cell numbers in the CA2 area of NTg mice housed in SH for 10 days. However, in Tg2576 mice, a persistent decrease in these cell numbers was observed in CA2, regardless of PBS or ChABC treatment, compared to NTg mice. *: p < 0.05, **: p < 0.01, *: p < 0.001, Two-way ANOVA with Tukey’s post hoc test**. (C)** Quantification of PV+ cells 10 days after EE shows that EE restores PV+ cell numbers in the CA2 area of Tg2576 mice, eliminating genotype differences. PNN disruption in the CA1 area with ChABC did not affect these numbers. **(D)** Cell counts are expressed as a percentage of the mean observed in Tg2576 mice housed in standard conditions and injected with PBS. A 10- day EE increased PV+/PNN+ and PV+ cell numbers in the CA2 area of Tg2576-PBS mice. ChABC injection slightly impacts this effect. *: p < 0.05; **: p < 0.01, One-way ANOVA with Dunnett’s post hoc test.

**Figure S2:**
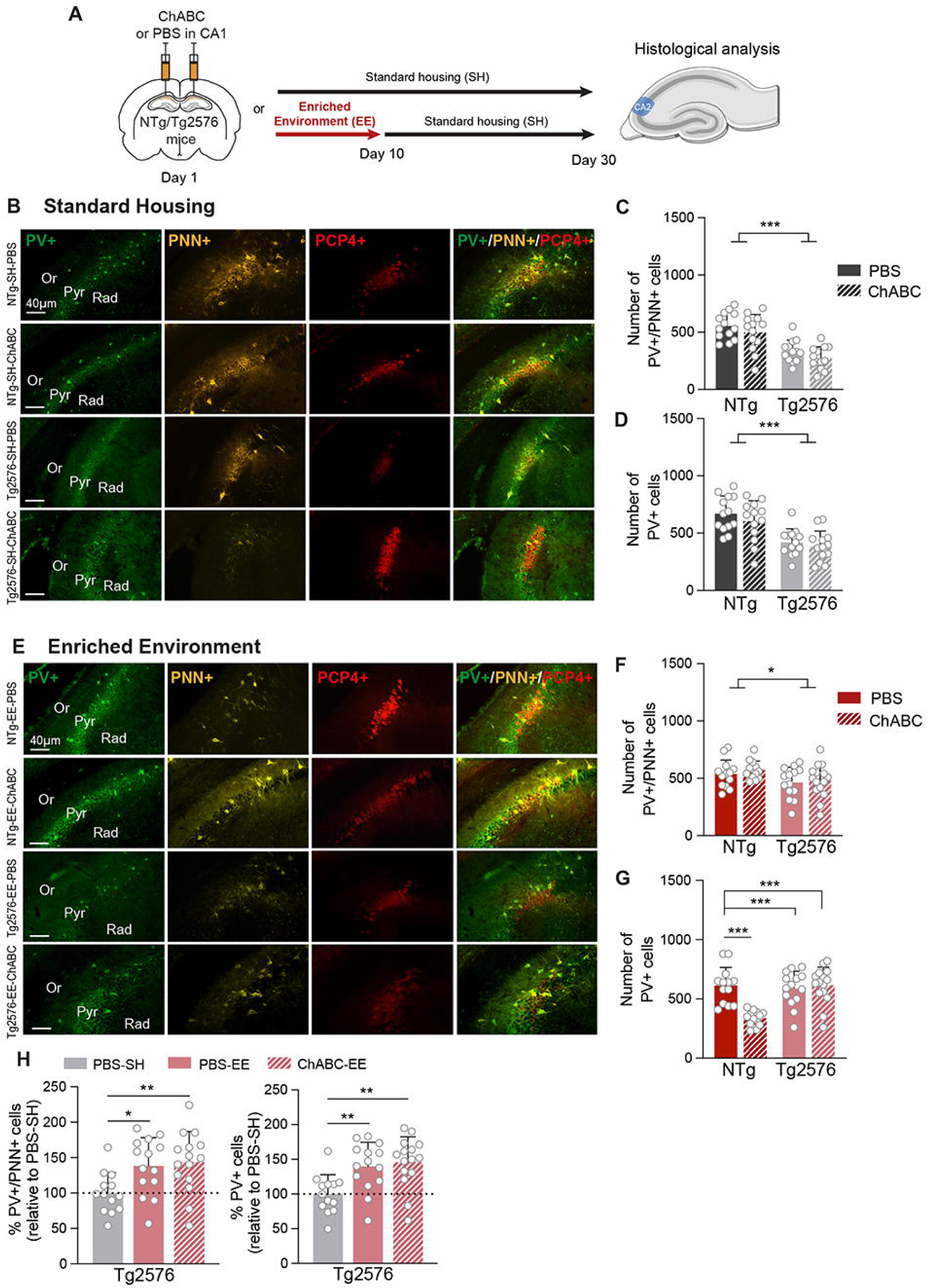
EE restore PV+/PNN+ cells number of CA2 area in Tg2576 mice at medium term. **(A)** Experimental timeline: ChABC or PBS was injected into the CA1 area of NTg and Tg2576 mice. Mice were either maintained in SH or placed in EE for 10 days before returning to SH. Thirty days after the injections, mice were sacrificed for anatomical quantification**. (B)** Microphotographs illustrating PV+ (green), PNN+ (WFA, yellow) and PCP4+ (red) cells in the CA2 area of NTg and Tg2576 mice injected with PBS or ChABC into the CA1 area, which were maintained in SH condition. **(C-D)** Quantification of PV+/PNN+ and total PV+ cells in CA2 shows a decrease in Tg2576 mice as compared to NTg mice housed in SH condition, regardless of treatment. *: p < 0.001, Two-way ANOVA with Tukey’s post hoc test. Bar graphs show means and SD. **(E)** Microphotographs illustrating PV+ (green), PNN+ (WFA, yellow) and PCP4+ (red) cells in the CA2 area of NTg and Tg2576 mice injected with PBS or ChABC into the CA1 area, and that spent 10 days in an EE. **(F-G)** Quantification of PV+/PNN+ and PV+ cells in the area CA2 of mice that spent 10 days in EE after injections reveals increased numbers in Tg2576 mice, regardless of PBS or ChABC injection into the CA1 area. Despite this increase, cell numbers in Tg2576 mice remain below those in NTg mice. *: p < 0.05; **: p < 0.01; *: p < 0.001, Two-way ANOVA with Tukey’s post hoc test. Bar graphs show means and SD. **(H)** Percentage of PV+/PNN+ and PV+ cell numbers relative to what was observed in Tg2576-PBS-SH mice across different housing and injection conditions. A 10-day EE increased PV+/PNN+ and total PV+ cell numbers in the CA2 area of Tg2576 mice, regardless of the nature of the injection into CA1. *: p < 0.05; : ** p < 0.01, One-way ANOVA with Dunnett’s post hoc test.

**Figure S3:**
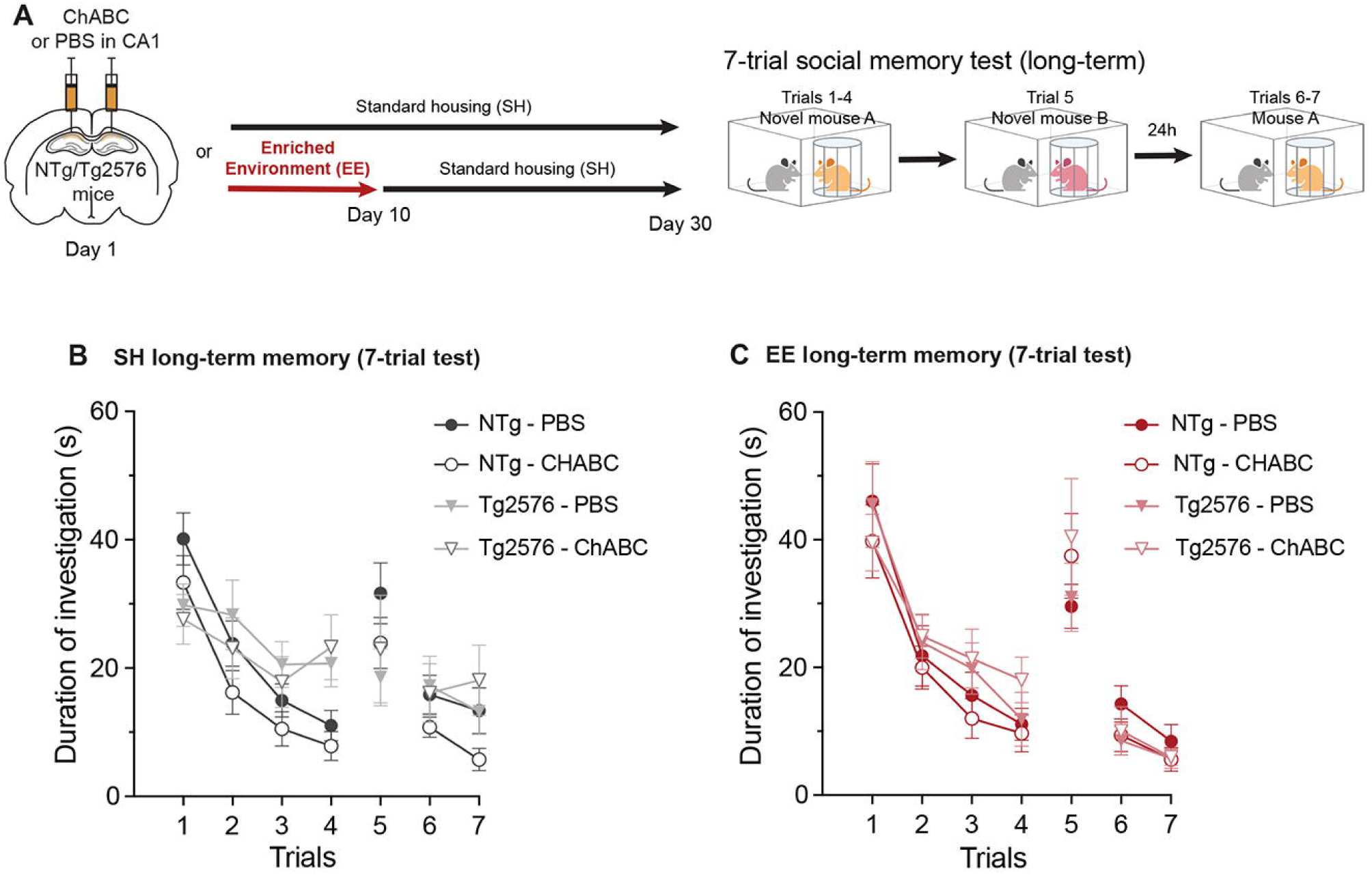
ChABC injection into the CA1 area does not impair EE-induced improvement of investigation time during social memory task in Tg2576 mice. **(A)** Animals were injected with PBS or ChABC into the CA1 area, then exposed to EE for 10 days followed by SH for 20 days, or remained in SH for 30 days. Afterward, mice were tested for spatial and social memory. **(B)** Total investigation time across the 7 trials shows that NTg mice habituate to the presented mouse during the first four trials and exhibit renewed interest in a new mouse at trial 5. On the next day, trials 6 and 7 demonstrate long-term retention for the first presented mouse by NTg mice. Tg2576 mice that stayed in SH, however, exhibit impaired social memory. **(C)** Following 10 days of EE, Tg2576 mice demonstrate a complete rescue of their social memory, as shown by learning curves across trials similar to those of NTg mice. Disrupting PNN in the CA1 area with ChABC injection did not dampen the beneficial effect of EE on social memory in Tg2576 mice.

